# Targeting *Bothrops asper* Venom Enzymes: Steroidal Derivatives as potential Inhibitors of Phospholipase A_2_, Serine proteinases, and metalloproteinases

**DOI:** 10.64898/2026.04.15.718639

**Authors:** Mitchell Bacho, Yeray A. Rodriguez-Núñez, Cristián J. Guerra, Efraín Polo-Cuadrado, Jorge Soto-Delgado, Ángela Sofia Tejada Restrepo, Julián Rodrigo Méndez Anacona, Isabel Henao-Castañeda, Lina María Preciado Rojo

## Abstract

Snakebite envenoming is a neglected tropical disease responsible for an estimated 1.8–2.7 million envenomings and 80,000–140,000 deaths annually, with *Bothrops asper* accounting for 66.7% of cases and 73.2% of deaths in Colombia. The inhibitory activity of three semi-synthetic ergosterol-derived compounds (**2**, **3**, and **4**) was evaluated against the major enzymes of *Bothrops asper* venom—snake venom metalloproteinases (SVMPs), phospholipases A₂ (PLA₂s), and serine proteinases (SVSPs)—through *in vitro* and *in silico* studies, aiming to identify potential adjuvants for the treatment of local damage. *In vitro* assays were developed to assess the inhibition of procoagulant, amidolytic, proteolytic, phospholipase A_2_, and esterase activities using compound concentrations ranging from 62.5 to 500 μM, along with molecular docking studies to predict enzyme-ligand interactions. Compound **4** was the most effective inhibitor of coagulant activity (SVSP), showing a significant dose-dependent effect (p < 0.001) at all tested concentrations (62.5–500 μM), prolonging plasma clotting time by up to 300 s at the highest doses. For amidolytic activity (SVSP), compounds **2** and **4** showed inhibitory capacity, although with variability across concentrations. In contrast, compounds **2** and **3** were the most potent inhibitors of PLA₂ activity inhibitors, exhibiting a significant dose-dependent effect. Notably, none of the compounds inhibited SVMP proteolytic activity. Molecular docking and molecular dynamics simulations were performed to investigate the binding mechanisms of the selected compounds with PLA_2_ and SVSPs. Compound **2** was analyzed in complex with PLA_2_, and compounds **3** and **4** were evaluated against SVSP. The results revealed that ligand binding was primarily driven by hydrophobic interactions, supported by key electrostatic contributions, leading to stable ligand–receptor complexes throughout the simulations. MM-GBSA calculations showed favorable binding free energies consistent with experimental inhibitory activity, highlighting ergostane-based compounds as promising scaffolds for the development of novel inhibitors targeting PLA_2_ and SVSP.

**Author summary:** Every year, hundreds of thousands of people are bitten by snakes, most of them farmers or children living in rural areas far from hospitals. Many suffer permanent damage or do not survive. Snakebite is a serious global health problem that rarely receives the attention it deserves.

In Colombia, *Bothrops asper* — known locally as *mapaná* or *terciopelo* — is responsible for most of these cases. Its venom acts quickly, destroying tissue, causing bleeding, and disrupting the blood’s ability to clot. Although treatments exist, they often cannot prevent the severe damage that occurs within the first minutes after a bite.

With this in mind, we explored whether molecules derived from ergosterol, a natural compound found in mushrooms, could help block some of the most harmful effects of the venom. Through laboratory experiments and computer simulations, we found that some of these molecules were able to reduce venom activity linked to tissue damage and clotting disorders, although they did not block all of its effects.

We hope these findings represent a step toward developing complementary treatments that are simpler and more accessible, ultimately improving care for the people who need it most.

## 1. Introduction

Snakebite envenoming is a neglected tropical disease that primarily affects rural communities in warm and humid regions, where access to specialized medical services is limited. In June 2017, the World Health Organization officially included it among the highest-priority tropical diseases, given its high morbidity, potential lethality, and limited availability of adequate treatments in vulnerable areas(1).

Each year, approximately 5.4 million snakebites are reported, of which 1.8-2.7 million results in envenomation, causing 80,000-140,000 deaths annually. Most cases occur in Africa, Asia, and Latin America. In Latin America, approximately 150,000 snakebite accidents occur each year, causing the death of approximately 5,000 people, with Brazil, Mexico, Venezuela, and Colombia being the most affected countries(2).

In Colombia, a country characterized by high biodiversity and venomous snakes at altitudes below 2,500 m.a.s.l., 16,570 confirmed snakebite cases were reported between 2022 and 2024. Snakes of the genus *Bothrops*, particularly *Bothrops asper*, are the primary cause of envenomation in the country, accounting for 66.7% of cases and 73.2% of associated deaths(3). These species, commonly known as *mapaná* or *terciopelo*, are widely distributed in the tropical regions of the Colombian Caribbean, Andes, and Pacific. Its defensive behavior when threatened makes it one of the most medically relevant species in Brazil(4). Most of these cases occur in rural areas and are associated with agricultural activities, walking along trails, or domestic tasks, with the departments of Antioquia, Córdoba, and Bolívar being the most affected(3).

Envenoming by *Bothrops asper* produces both local and systemic clinical manifestations. Local effects included edema in 95% of cases, detectable within 5 min after the bite, local hemorrhage in 34% of cases, and tissue necrosis in 10% of cases. Systemic effects may include defibrinogenation, thrombocytopenia, hypotension, and internal hemorrhage(4).

These symptoms are due to the action of various enzymes present in the venom, among which metalloproteinases (SVMP), phospholipases A_2_ (PLA_2_), and thrombin-like serine proteinases (SVSP) are prominent. These enzymes are responsible for myotoxic, hemotoxic, and coagulation-disrupting effects, causing tissue and systemic damage in envenomed patients(5).

Conventional treatment for these accidents consists of intravenous administration of antivenom obtained from immunized animals, such as horses and sheep(6). Although effective in counteracting systemic effects, these antivenoms do not remedy the rapidly developing local effects caused by major venom toxins if not administered immediately (7). Moreover, their application can trigger adverse reactions due to nonspecific immune responses (8).

Given these limitations, there has been growing interest in the search for alternative neutralizing agents from diverse natural and synthetic sources to develop new therapeutic tools that complement and improve conventional antivenom therapy (9). Among these advances, two ergosterol-derived compounds have been reported to inhibit phospholipases A_2_ (PLA_2_) from the venom of *Crotalus adamanteus*(10), which belongs to the Viperidae family. Although venom composition varies greatly between species and even within the same species depending on environmental factors, age, sex, or prey availability(11), this finding suggests the potential of ergosterol derivatives for therapeutic use. Based on the aforementioned, the present study evaluated the inhibitory activity of three synthetic ergosterol-derived compounds against the major enzymes of *B. asper* venom using in vitro and in silico approaches, with the aim of contributing to the research and development of potential adjuvants for the treatment of local damage caused by snakebite.

## 2. Experimental

### 2.1. Material and methods

The solvents were dried and distilled using standard methods before use. The reagents were purchased from commercial suppliers and purified further. Ergosterol was purchased from Cayman Chemical (Ann Arbor, MI, USA). ^1^H and ^13^C NMR spectra were recorded on a Bruker Advance III HD-400 spectrometer by dissolving the samples in deuterated chloroform (CDCl_3_). Chemical shifts (*δ*) are reported in ppm, relative to TMS as an internal standard. The coupling constants (J) are expressed in Hertz (Hz), and the integrations are reported as the number of protons. The following abbreviations are used to describe the peak patterns: s for singlet, d for doublet, t for triplet, m for multiplet, and br for broad. Two-dimensional spectra were obtained using standard Bruker software.

### 2.2. Synthesis and characterization of ergosterol derivatives

#### 2.2.1. Synthesis of 4-phenyl-1,2,4-triazoline-3,5-dione (1)

The synthesis of the SiO₂-HNO₃ mixture and subsequent 4-phenylurazole oxidation were carried out using the method proposed by Ghorbani(12). In a round-bottom flask equipped with a stirring bar, 2.82 g of HNO₃ (65% v/v) was added to 2.0 g of SiO₂, and the mixture was stirred for 10 min at room temperature. A white solid containing the SiO₂-HNO₃ mixture was obtained. Subsequently, 4-phenylurazole was oxidized by dissolving 3.544 g of 4-phenylurazole (20 mmol) in 100 mL of CH₂Cl₂ and gradually adding 4.0 g of the SiO ₂-HNO ₃ mixture. After stirring the solution for 30 min, anhydrous sodium sulfate was added, and the mixture was filtered after 20 min. The solvent was evaporated under reduced pressure, yielding a pink solid corresponding to 4-phenyl-1,2,4-triazoline-3,5-dione with a yield of 95% (3.33 g).

#### 2.2.2. Synthesis of PTAD-protected ergosterol (3)

To a round-bottom flask equipped with a stirring bar, 3.97 g of ergosterol (**2**, 10.0 mmol) was added and dissolved in 168 mL of acetone. The mixture was stirred while a solution of 1.93 g of 4-phenyl-1,2,4-triazoline-3,5-dione (**1**, 11.0 mmol) in 94 mL of acetone was added dropwise using an addition funnel for 15 min. Before the addition, TLC monitoring confirmed the complete consumption of ergosterol, the mixture was concentrated in vacuo, and the crude product was purified using a silica gel column with a hexane/ethyl acetate solution (1:1 v/v), yielding compound 1 as a light-yellow solid (5.18 g, 90.0%). R_f_ value was 0.29 (ethyl acetate:hexane, 1:1 v/v). ¹H NMR (400 MHz, CDCl₃) δ ppm: 7.46 – 7.35 (m, 4H), 7.33 – 7.27 (m, 1H), 6.36 (d, *J* = 8.3 Hz, 1H), 6.19 (d, *J* = 8.3 Hz, 1H), 5.28 – 5.12 (m, 2H), 4.39 (tt, *J* = 10.6, 5.3 Hz, 1H), 3.13 (dd, J = 14.1, 4.8 Hz, 1H), 2.91 (s, 1H), 1.03 (d, *J* = 6.6 Hz, 3H), 0.92 (s, 3H), 0.90 (d, *J* = 6.8 Hz, 3H), 0.83 (d, *J* = 6.8 Hz, 3H), 0.80 (s, 3H). ^13^C NMR (100 MHz, CDCl_3_) δ ppm: 148.81 (C=O), 146.10 (C=O), 135.69 (CH=C), 135.09 (CH=C), 132.39 (CH=C), 131.75 (C=C), 129.02 (CH=C), 128.84 (CH=C), 127.78 (CH=C), 126.17 (CH=C), 67.25 (C), 65.80 (CH), 64.95 (C), 55.14 (CH), 52.90 (CH), 49.29 (CH), 43.86 (C), 42.78 (CH), 41.19 (CH_2_), 39.53 (CH), 38.14 (C), 34.71 (CH), 34.20 (CH), 33.06 (CH_2_), 29.00 (CH_2_), 27.53 (CH_2_), 23.35 (CH_2_), 22.42 (CH_2_), 21.24 (CH_3_), 19.95 (CH_3_), 19.63 (CH_3_), 17.62 (CH_3_), 17.55 (CH_3_), 13.25 (CH_3_).

#### 2.2.3. Oxidation of PTAD-ergosterol to PTAD-ergosta-3-one (4)

In a round-bottom flask equipped with a stirring bar, PTAD-ergosterol (**3**, 4.57 g, 8.0 mmol), calcium carbonate (CaCO₃, 5.16 g, 27 mmol), and 4 Å molecular sieves (2.823 g) were stirred in 283 mL of dichloromethane. Then, pyridinium chlorochromate (PCC, 5.82 g, 27 mmol) was added, and the slurry was stirred overnight at room temperature. To remove the generated chromium salts, 300 mL of diethyl ether was added to the reaction mixture, which was then passed through a Celite 545 and silica gel pad. The solution obtained was concentrated under reduced pressure to yield PTAD-protected ergosta-3-one as a light-yellow solid (4.87 g, 94.5% yield). R_f_ value was 0.36 (ethyl acetate: hexane, 1:1 v/v). ¹H NMR (400 MHz, CDCl₃) δ ppm: 7.48 (m, 5H), 7.38 (m, 1H), 6.64 (d, *J* = 9.5 Hz, 1H), 6.55 (d, *J* = 8.3 Hz, 1H), 5.36 – 5.14 (m, 3H), 3.58 (d, *J* = 18.3 Hz, 1H), 2.79 (d, *J* = 18.3 Hz, 1H), 1.21 (s, 3H), 1.06 (s, 2H), 0.91 (d, *J* = 6.9 Hz, 4H), 0.87 (s, 2H), 0.83 (d, *J* = 6.6 Hz, 7H), 0.81 (s, 2H), 0.77 (s, 3H). ^13^C NMR (100 MHz, CDCl_3_) δ ppm: 204.40 (C=O),149.51 (C=O), 146.81 (C=O), 135.65 (CH=C), 135.11 (CH=C), 132.59 (CH=C), 131.41 (C=C), 129.26 (CH=C), 129.03 (CH=C), 128.10 (CH=C), 126.01 (CH=C), 65.51 (CH), 64.19 (C), 55.26 (CH), 53.95 (CH), 49.06 (CH), 44.04 (C), 42.89 (CH), 42.11 (CH_2_), 39.74 (CH), 38.61 (C), 34.84 (CH), 33.69 (CH), 33.18 (CH_2_), 28.41 (CH_2_), 28.07 (CH_2_), 23.77 (CH_2_), 22.81 (CH_2_), 21.07 (CH_3_), 20.07 (CH_3_), 19.78 (CH_3_), 17.68 (CH_3_), 17.57(CH_3_), 13.18 (CH_3_).

### 2.3. Venoms

A venom pool was obtained by manual extraction from *adult B.* asper specimens from the subregions of Antioquia, Magdalena Medio, and Meta in the southeast region of Colombia. The snakes were maintained in captivity at the Serpentarium of the Universidad de Antioquia (Medellín, Colombia). The venoms were centrifuged at 5000 rpm for 15 min, and the supernatants were lyophilized and stored at −20 °C until use.

### 2.4. *In vitro* procoagulant activity of Serine proteinases (SVSP)

The procoagulant activity of the venom in human citrated plasma was determined using the modified method described by Theakston and Reid (13). The minimum coagulant dose (MCD) was set at 1 μg of *B. asper* venom (a dose that induces blood plasma coagulation in 20 s), diluted 1:1 in saline solution. For the assay, venom mixtures at this dose were prepared with four different concentrations of ergosterol-derived compounds (62.5, 125, 250, and 500 μM). The mixtures were preincubated at 37 °C for 30 min, after which 100 μL of each treatment was added to 200 μL of citrated human plasma. The plasma clotting time at 37 °C was recorded, with a maximum observation time of 30 min. Each assay was performed in quadruplicate. Venom was used as the positive control, and the compounds at the highest concentration tested served as the negative control (14).

### 2.5. Inhibition Assay of the Amidolytic Activity of SVSP

BAPNA (*N*-succinyl-arginine-*p*-nitroanilide, Sigma-Aldrich) was used to evaluate the amidolytic activity of SVSP and its inhibition by ergosterol-derived compounds. For this, mixtures of *B. asper* venom (3 μg/μL) with four different concentrations of the derivatives (62.5, 125, 250, and 500 μM) were prepared and pre-incubated at 37 °C for 30 min. Subsequently, 10 μL of each treatment was added to 200 μL of the substrate (10 mM) with 50 μL of buffer. The mixtures were incubated for 40 min at 37 °C, and the absorbance was measured at 405 nm. Each assay was performed in quadruplicate. Venom was used as the positive control, and the compounds at the highest concentration tested were used as the negative control(15).

### 2.6. Inhibition Assay of the Proteolytic Activity of SVMP

Proteolytic activity was evaluated using azocasein as a substrate, as described by Wang et al. The minimum proteolytic dose (MPD) was set at 20 μg of *B. asper* venom (a dose that produces a 0.5 change in absorbance relative to the negative control), diluted in 20 μL of buffer (25 mM Tris, 150 mM NaCl, 5 mM CaCl₂, pH 7.4). For the assay, mixtures of venom at this dose with four different concentrations of ergosterol-derived compounds (62.5, 125, 250, and 500 μM) were prepared. The mixtures were pre-incubated at 37 °C for 30 min. Subsequently, 100 μL of azocasein (10 mg/mL) was added, and the mixture was incubated for 90 min at 37 °C. The reaction was stopped by adding 200 μL of 5% trichloroacetic acid, and the samples were centrifuged at 3000 rpm for 5 min to separate the precipitate. Finally, 100 μL of the supernatant was collected and mixed with 100 μL of 0.5 M NaOH solution. Each assay was performed in quadruplicate, and the absorbance was measured at 450 nm using a spectrophotometer. Venom was used as the positive control, and the compounds at the highest concentration tested were used as the negative control(16,17).

### 2.7. Inhibition Assay of the Esterase Activity of SVPLA₂s

The inhibition of PLA₂ esterase activity was evaluated by the cleavage of the synthetic substrate 4-NOBA (4-nitro-3-(octanoyloxy) benzoic acid). Mixtures of *B. asper* venom with four different concentrations of ergosterol-derived compounds (62.5, 125, 250, and 500 μM) were prepared. The mixtures were pre-incubated at 37 °C for 30 min. Subsequently, 25 μL of these treatments were added to 200 μL of buffer (10 mM Tris-HCl, 10 mM CaCl₂, 100 mM NaCl, pH 8.0) and 25 μL of 4-NOBA (1 mg/mL). The final reaction mixture was incubated at 37 °C for 60 min, after which the absorbance was measured at 425 nm. Each assay was performed in quadruplicate. Venom was used as the positive control, and the compounds at the highest concentration tested were used as the negative control(18).

### 2.8. Inhibition of phospholipase A_2_ Activity Using EnzChek™ PLA_2_ Assay Kit

PLA_2_ activity was estimated using the EnzChek™ PLA_2_ assay kit (Thermo Fisher Scientific, Waltham, MA, USA) following the manufacturer’s protocol. A liposome mix was prepared by adding 30 μL each of 10 mM dioleoyl phosphatidylcholine (DOPC), 10 mM dioleoyl phosphatidylglycerol (DOPG), and red/green BODIPY® PC-A2. Liposomes used as PLA_2_ substrates were formed by slowly adding 90 µL of the lipid mixture to 9 mL PLA_2_ buffer. Two micrograms of *B. asper* venom was used to estimate PLA_2_ activity (positive control). The samples for the positive control, venoms, and ergosterol derivatives (50 µL) were added to 96-well opaque plates along with 50 µL PLA_2_ substrate, and the relative fluorescence units (RFU) were recorded after 10 min of incubation in a Fluoroskan Ascent (Thermo Scientific) with an emission wavelength of 515 nm and an excitation wavelength of 480 nm. The experiment was performed in triplicate.

### 2.9. Statistical Analysis

A two-way ANOVA was used to compare the positive controls with the results of the different mixtures of venom and ergosterol-derived compounds in each in vitro assay. Subsequently, a Bonferroni post hoc test was performed to assess the differences between the means. Differences were considered significant at *p* < 0.05.

### 2.10. Molecular Docking

Automated docking was employed to determine the optimal binding orientations and conformations of compound **2** on PLA_2_ and compounds **3** and **4** within the SVSP binding site. The crystal structure of PLA_2_ used in the molecular docking simulations was obtained from the RCSB Protein Data Bank with PDB ID 5TFV.

SVSP from *B. asper* venom was modeled and evaluated (Q072L6 · VSPL_BOTAS). The sequence was obtained from UniProt(19). The signal peptide and propeptide were manually removed. The resulting FASTA sequence was submitted to the AlphaFold server, which uses AlphaFold3. In addition, the stereochemical quality of the modeled protein structure and its overall structural geometry were confirmed using PROCHECK(20).

Molecular docking calculations were performed using AutoDock 4.2 software(21) and PyMOL version 2.5 Molecular Graphics System(22) with the AutoDock/Vina plugin(23). The protein structure was prepared by removing water molecules and incorporating the Kollman charges and polar hydrogen atoms. After docking, the docked poses were clustered into groups with RMSD values below 1.0 Å. From the ensemble of predicted molecular complexes, the most populated cluster conformation and the lowest-energy conformation for the most active compound docked to the receptor were selected. Following molecular docking calculations, all protein-ligand complexes were prepared in PDB format using the Protein-Ligand Interaction Profiler (PLIP) version 2.4 software(24) and subsequently visualized using PyMOL version 2.5 (22).

### 2.11. Molecular dynamics simulations

The structures derived from molecular docking calculations were further studied using MD simulations to obtain energy data for the ligand-receptor complex and the interactions among the residues involved. For MD simulations, the best-posed structures were selected based on the molecular docking results. MD simulations were performed using Amber 22 software (25). A ligand topology builder (ACPYPE) was used to generate the parameters for all ligands(26,27). GAFF2, ff19SB, and TIP3P force fields were used to describe the compounds, proteins, and water, respectively. Solvated complex minimization, heating to 300 K for 1 ns, density equilibration for 1 ns, and constant pressure equilibration for 5 ns were performed. The production phase of 500 ns was run, and the coordinates were recorded every 10 ps. Simulations were performed with a 2 fs time step and Langevin dynamics for temperature control. MD trajectories were automatically clustered using TTClust version 4.7.2 and the elbow method to calculate the number of clusters, and a representative structure for each cluster was produced(28). Representative structures were analyzed using PLIP version 2.4 to determine the types and numbers of bonds participating in the interactions [24]. Additionally, the interaction energy for **2-4**, expressed as ΔG_bind_, was estimated using the Molecular Mechanics Generalized Born Surface Area (MM/GBSA) method with the MMPBSA.py.MPI module implemented in Amber 22 to compare the most favorable interactions of the inhibitors(29,30). The three-dimensional structures of the protein-ligand complexes and MD trajectory analysis were visually inspected using the computer graphics programs VMD version 1.93 (30) and PyMOL version 2.5 (22).

## 3. Results and discussion

### 3.1. Synthesis and characterization of compound 1-3

The synthetic route for the formation of ergosterol derivatives **2-3** is described in Figure 1. The modification of 4-phenylurazole to 4-phenyl-1,2,4-triazoline-3,5-dione can be performed using homogeneous and heterogeneous catalysis(31–34). In our case, the use of SiO₂/HNO₃ as an oxidant mixture successfully yielded 4-phenyl-1,2,4-triazoline-3,5-dione **2** with a high yield (95%). This intermediate was used immediately in the next synthetic step. Compound **3** was obtained via a [4+2] Diels-Alder cycloaddition reaction between **2** and ergosterol, both dissolved in acetone. The structure of this intermediate was determined using NMR spectroscopy. The ¹H NMR spectrum of **2** showed five signals associated with methyl groups in the range of 0.80–1.03 ppm, indicating that the compound is a derivative of the ergostane skeleton. Additionally, at δ_H_ 4.39 ppm (m, 1H, H_3_), the hydroxyl group present in ring A can be identified, which has not been oxidized. Due to the unsaturation present in ergosterol, two signals associated with the alkenes in the side chain and B ring were identified. The signal at δ_H_ 5.20 ppm (m, 2H, H_22_-H_23_) corresponds to the hydrogens at positions C22 and C23 of the side chain, exhibiting a coupling constant of 15.2 Hz, consistent with trans isomerism of the alkene. Likewise, the signals at δ_H_ 6.19 ppm and 6.39 ppm (d, *J*=8.3 Hz, 1H, H6-H7) confirm the correct [4+2] cycloaddition between ergosterol and compound **1**. For this fragment, distinctive signals were detected at δ_H_ 7.29 ppm (m, 1H) and 7.41 ppm (m, 4H), indicating the presence of a phenyl group originating from the 4-phenyltriazolidone fragment. This information confirmed the successful incorporation of compound **1** into ergosterol.

**Figure 1.**
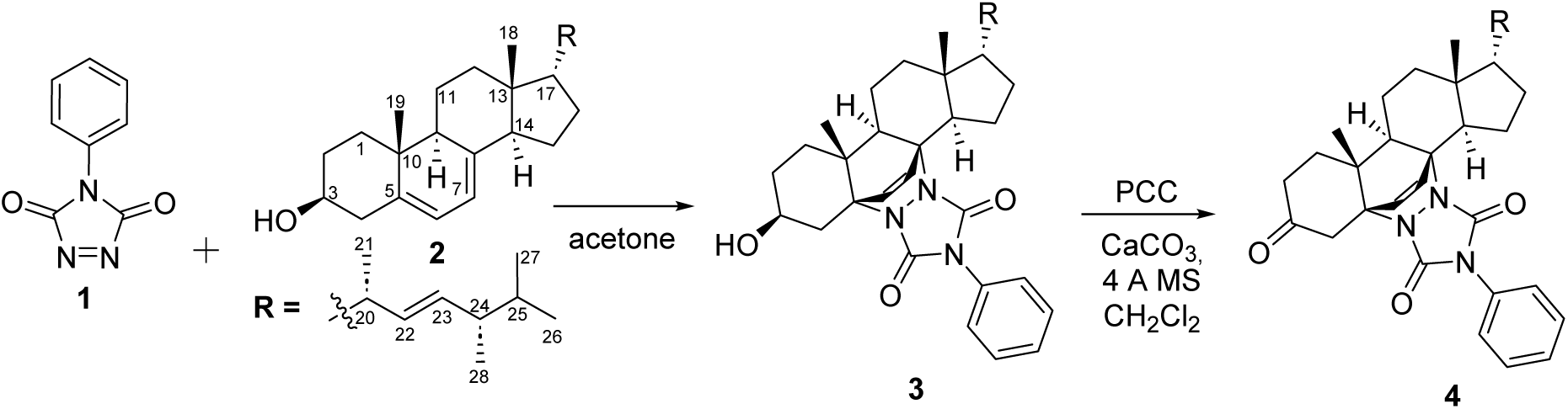
Synthetic scheme for the synthesis of PTAD-ergostanes (**3-4**) derivates from ergosterol.

Similarly, the oxidation of compound **2** with PCC, followed by chromatographic separation and subsequent NMR characterization, showed a pattern associated with the six methyl groups in the ergostane ring, between δ_H_ 0.77 and 1.06 ppm. Additionally, the absence of the signal at δ_H_ 4.18 ppm (m, 1H), corresponding to H3, and the presence of a δ_C_ 204.4 ppm (C3) signal confirmed the formation of the endocyclic ketone. In parallel, signals associated with the 4-phenyltriazolidinone fragment present in the B ring of the ergostane skeleton were observed at δ_H_ 7.30 ppm (1H, m) and 7.38 ppm (4H, m). This evidence supports the successful oxidation of PTAD-ergosta-3-one **3**.

### 3.2. Venoms

*Bothrops asper* is the species with the greatest clinical relevance in Colombia and other regions(35). Its venom is mainly composed of proteins and peptides, with the most abundant components being metalloproteinases (SVMPs) (33.2%), phospholipases A ₂ (PLA₂s) (31.3%), and serine proteinases (SVSPs) (9.3%) (36). These enzymes are the main toxic constituents of the venom, as they induce edema, myotoxicity, hemorrhage, and coagulation disorders, among other effects(11). Consequently, the ability of ergosterol-derived compounds to inhibit enzymatic and biological activities induced by these toxins was evaluated.

### 3.3. Inhibition of Procoagulant Activity

Compound **4** exhibited the greatest ability to inhibit venom-induced procoagulant activity in vitro. This inhibition was dose-dependent, with statistically significant differences relative to the positive control at all concentrations tested (62.5–500 μM; p < 0.001). The positive control, corresponding to untreated venom, showed a clotting time of approximately 20 s. In contrast, compounds **2** and **3** inhibited procoagulant activity only at the highest concentration tested (500 μM), with no significant effects at lower concentrations (Figure 2).

**Figure 2.**
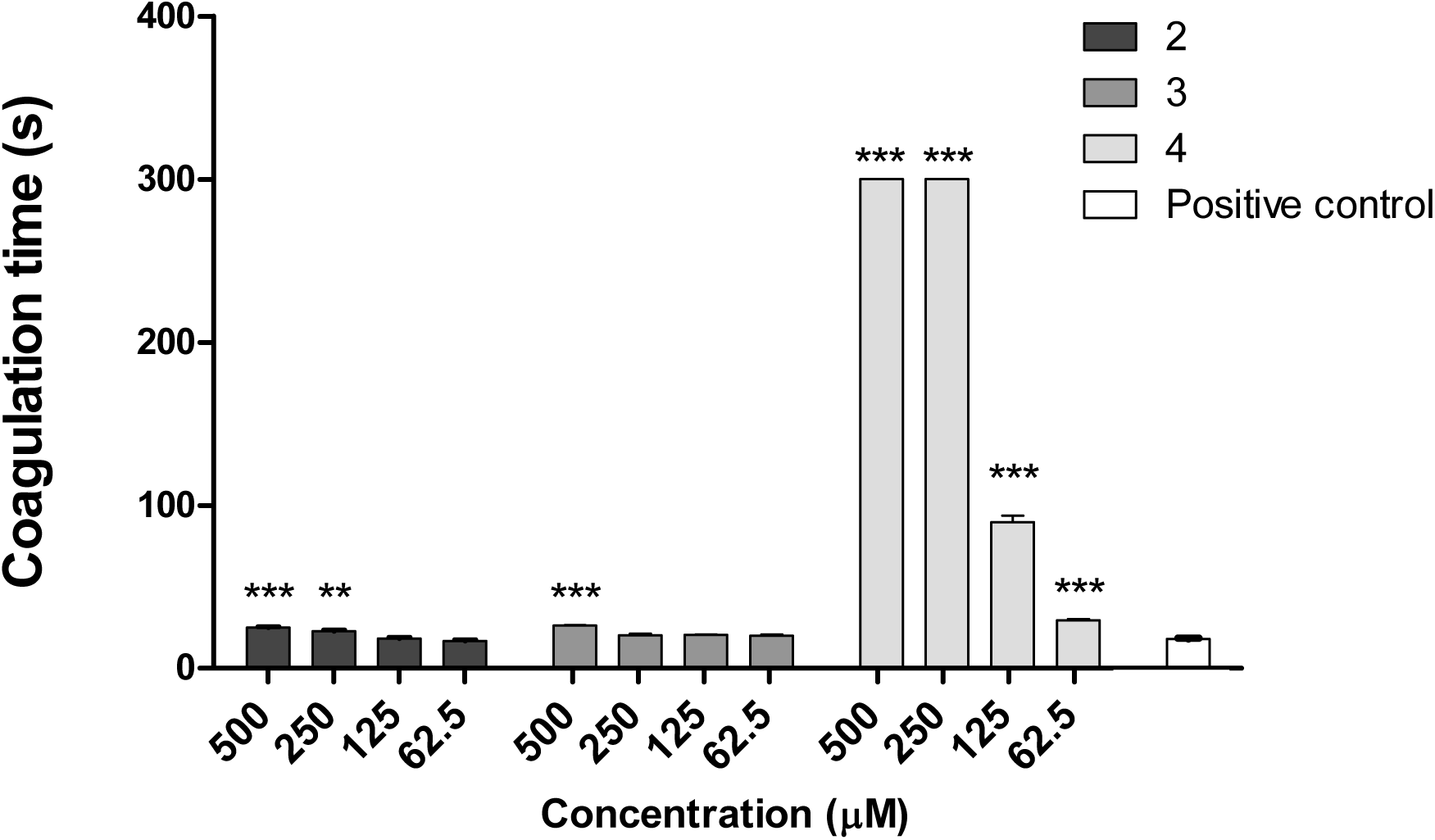
Inhibition of coagulant activity by ergosterol and ergostane-derived compounds. Data are presented as mean clotting time ± SEM (n = 3). *** indicates statistically significant differences compared with the positive control at p < 0.001, ** at p < 0.01, and * at p < 0.05.

SVSPs are among the main protein families implicated in alterations in human hemostasis, specifically thrombin-like enzymes (TLEs), which recognize and cleave human fibrinogen, releasing fibrinopeptide A or B, and do not act on coagulation Factor XIII. These toxins contribute to coagulopathy by consuming fibrinogen, a major systemic hemostatic disturbance frequently observed in snakebite victims. Thus, the procoagulant in vitro activity of *B. asper* venom correlates with its anticoagulant effect *in vivo* (37). These results demonstrate the potential of compound **4** to inhibit the anticoagulant effect induced by *Bothrops* envenomation; however, this must be confirmed in vivo studies.

### 3.4. Inhibition of Amidolytic Activity

The chromogenic substrate BAPNA was used to measure the amidolytic activity of *B. asper* venom, as it is hydrolyzed by snake venom serine proteases after the arginyl residue. Compounds **2** and **4** exhibited the highest inhibitory capacity against the amidolytic activity induced by *500 µM B. asper venom*. Compound **3** inhibited enzymatic activity at 250 µM but not at 500 µM (Figure 3).

**Figure 3.**
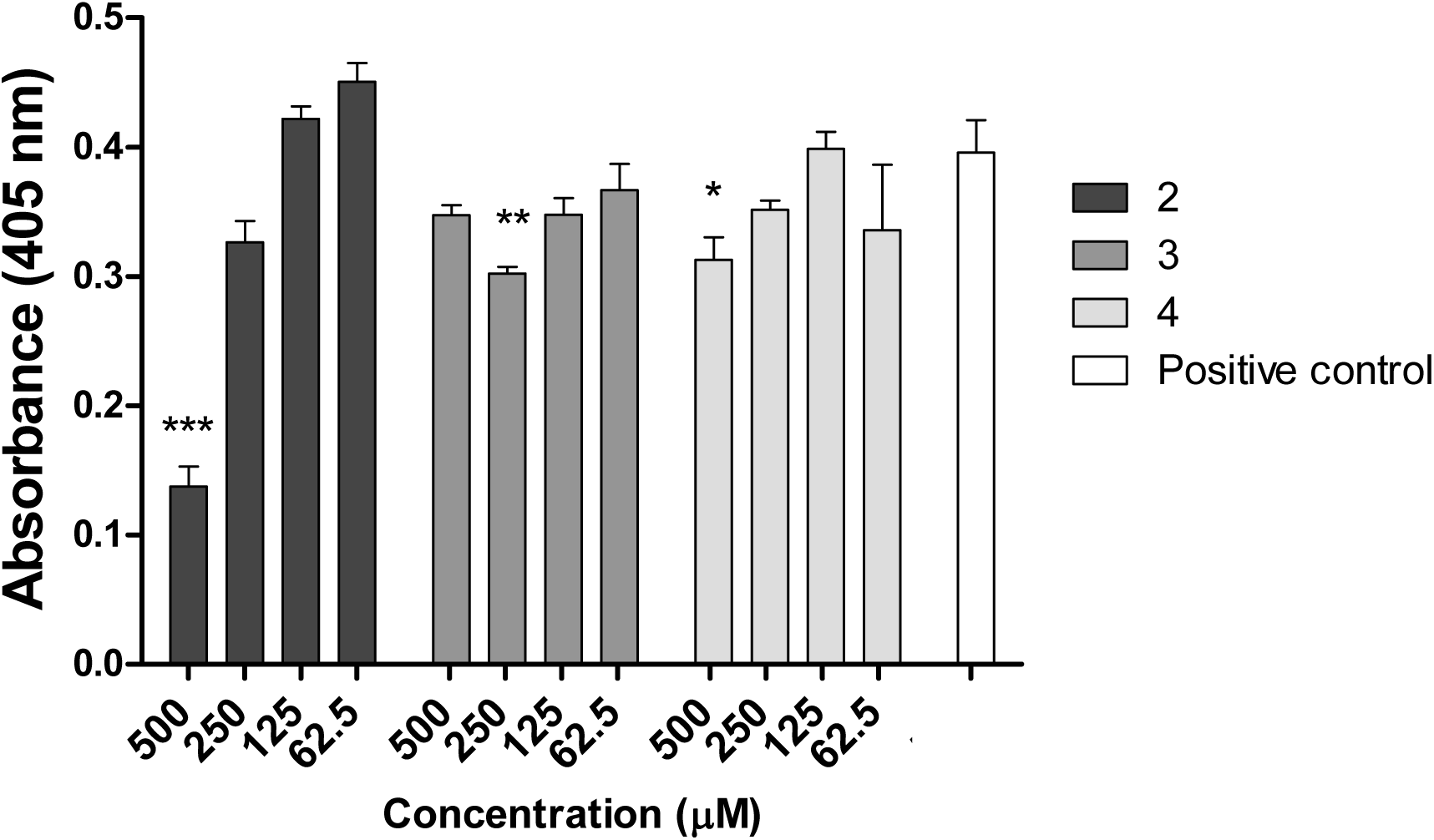
Inhibition of amidolytic activity by ergosterol and ergostane-derived compounds. Data are presented as mean clotting time ± SEM (*n* = 3). *** indicates statistically significant differences compared with the positive control at *p* < 0.001, ** at *p* < 0.01, and * at *p* < 0.05.

The inhibitory profile observed for compound 4 suggests that the procoagulant activity of *Bothrops asper* venom cannot be attributed exclusively to thrombin-like SVSPs. Although these enzymes play a central role in fibrinogen cleavage and consumption coagulopathy, other toxin families also significantly contribute to venom-induced hemostatic disturbances. Among these, phospholipases A₂ (PLA₂s) have been shown to exert both anticoagulant and procoagulant effects, depending on their structural features and interactions with coagulation complexes. Certain PLA₂ isoforms can indirectly enhance coagulation by promoting the availability of phospholipid surfaces, facilitating the assembly of prothrombinase and tenase complexes, or modulating membrane dynamics. Additionally, metalloproteinases (SVMPs) and C-type lectin-related proteins may also influence clot formation by targeting coagulation factors(38). Therefore, the significant inhibition of procoagulant activity by compound 4, along with its effect on amidolytic activity, suggests a broader mechanism of action potentially involving multiple enzymatic targets beyond SVSPs. This multitarget inhibition could be particularly relevant in the context of *Bothrops* envenomation, where synergistic interactions among toxin families determine the overall hemostatic outcome of envenomation. Further studies are required to elucidate the relative contributions of these enzymatic components and to confirm whether compound 4 directly interacts with PLA₂s or other non-serine protease toxins.

### 3.5. Inhibition of PLA₂ Activity

#### 3.5.1. PLA₂ Activity upon 4-NOBA substrate

The activity of PLA₂s present in snake venoms is implicated in phenomena such as myotoxicity, edema formation, and neurotoxicity, among other clinical manifestations(39). Compound **2** exhibited the greatest ability to inhibit PLA₂ activity induced by *B. asper* venom, using 4-NOBA as the substrate. This inhibition was concentration-dependent, with statistically significant differences compared to the positive control at all concentrations tested (62.5–500 μM; *p* < 0.001). Compounds **3** and **4** also exhibited inhibitory activity, albeit to a lesser extent (Figure 4).

**Figure 4.**
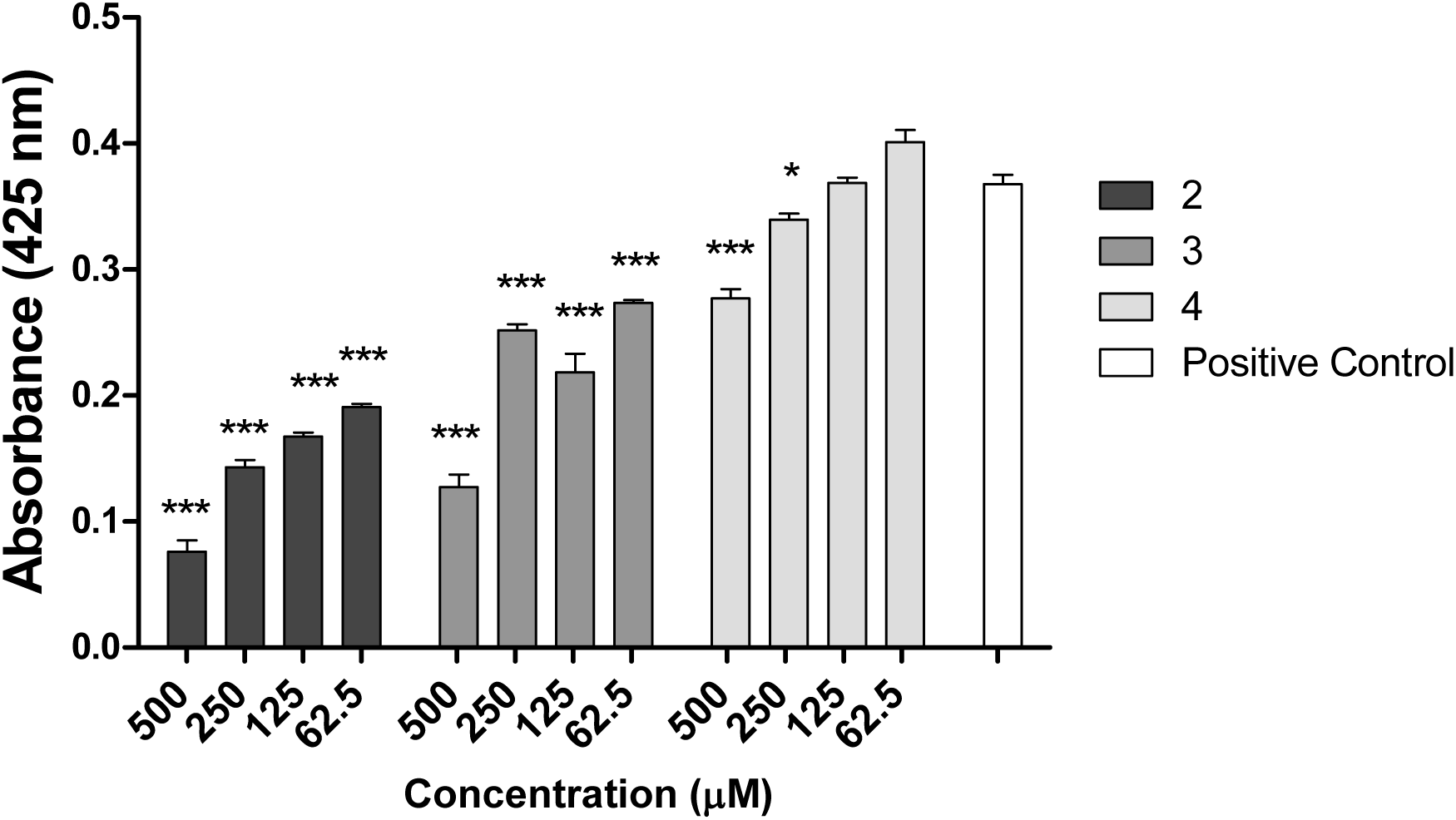
Inhibition of *B. asper* venom PLA₂ activity by ergosterol and ergostane-derived compounds using the monodisperse substrate 4-NOBA. Data are presented as absorbance ± SEM (n = 3). *** indicates statistically significant differences compared with the positive control at *p* < 0.001, ** at *p* < 0.01, and * at *p* < 0.05.

Previous research with ergosterol derivatives isolated from the fruiting bodies of a basidiomycete fungus, *Lactarius hatsudake*, 5α,8α-epidioxy-(22E,24R)-ergosta-6,22-dien-3β-ol and 5α,8α-epidioxy-(24S)-ergosta-6-en-3β-ol showed inhibitory activity against *Crotalus adamanteus* venom PLA_2_ enzyme(40). These results demonstrate the potential of ergosterol derivatives to inhibit these enzymes.

#### 3.5.2. Inhibition of phospholipase A_2_ Activity Using a fluorescent substrate

PLA_2s_ are a large class of interfacially active enzymes that catalyze the hydrolysis of glycerophospholipids at the sn-2 position. PLA_2s_ are weakly active on monomeric substrates such as 4-NOBA but are very active on organized types of substrates such as micelles, liposomes, or bilayers; therefore, it is not surprising that the enzyme activity is controlled by the physical properties of the substrate to a certain extent (41). Therefore, a liposomal substrate was chosen for the inhibition assays, as it more closely mimics the natural membrane environment and provides a more physiologically relevant platform for evaluating enzymatic activity, enabling an accurate assessment of inhibitory effects under conditions that closely resemble the native interfacial context of PLA_2_ action. In this assay, compound **3** exhibited the greatest ability to inhibit PLA₂ activity induced by *B. asper* venom across all evaluated concentrations. This inhibition was concentration-dependent, with statistically significant differences compared to the positive control at all concentrations tested (62.5–500 μM; *p* < 0.001). Compounds **2** and **4** also exhibited inhibitory activity at 500 μM (Figure 4).

The differential inhibitory profiles of compounds 2–4 in both PLA₂ assays reveal the complexity of targeting snake venom phospholipases and highlight the importance of substrate organization in modulating enzyme activity. While compound 2 exhibited the strongest inhibition in the 4-NOBA assay, compound 3 displayed superior inhibitory activity in the liposome-based system, suggesting distinct modes of interaction with PLA₂ enzymes depending on the physicochemical context of the substrate.

Secreted phospholipases A₂ (sPLA₂s) from viperid venoms, such as those present in *Bothrops asper*, are interfacially activated enzymes whose catalytic efficiency is markedly enhanced when acting on aggregated phospholipid substrates, including membranes and lipoprotein-like structures(42). In this regard, the limited activity observed with monomeric substrates, such as 4-NOBA, reflects only the intrinsic catalytic potential, whereas liposomal systems more accurately reproduce the interfacial activation process that occurs in vivo. This distinction is critical because inhibitors may differentially affect catalytic residues, calcium-binding loops, or membrane-docking regions, leading to variable efficacy across assays.

The higher inhibitory activity of compound 3 in the liposomal assay suggests that it may interfere with interfacial binding or disrupt enzyme–membrane interactions rather than directly targeting the catalytic site. Conversely, the stronger inhibition by compound 2 in the 4-NOBA assay may indicate a more direct interaction with the catalytic machinery of PLA₂, possibly involving interference with the His-Asp catalytic dyad or calcium-dependent stabilization of the active site.

Taken together, these results show that the evaluation of PLA₂ inhibitors should incorporate both monomeric and interfacial substrates to comprehensively capture their inhibitory potential and mechanism of action. The different behaviors of compounds 2 and 3 further suggest that combining inhibitors with complementary mechanisms could enhance the neutralization of PLA₂-driven toxicity in Bothrops envenomation. Future studies should focus on binding assays and structural approaches to characterize the precise molecular interactions involved and validate these findings *in vivo*.

### 3.6. Inhibition of proteolytic activity

None of the three ergosterol-derived compounds evaluated (**2**, **3**, and **4**) inhibited the proteolytic activity associated with SVMPs. No statistically significant differences were observed compared to the positive control at any of the tested concentrations (Figure 5). This lack of inhibition suggests that these compounds do not effectively interact with the catalytic machinery or structural domains required for SVMP activity.

**Figure 5.**
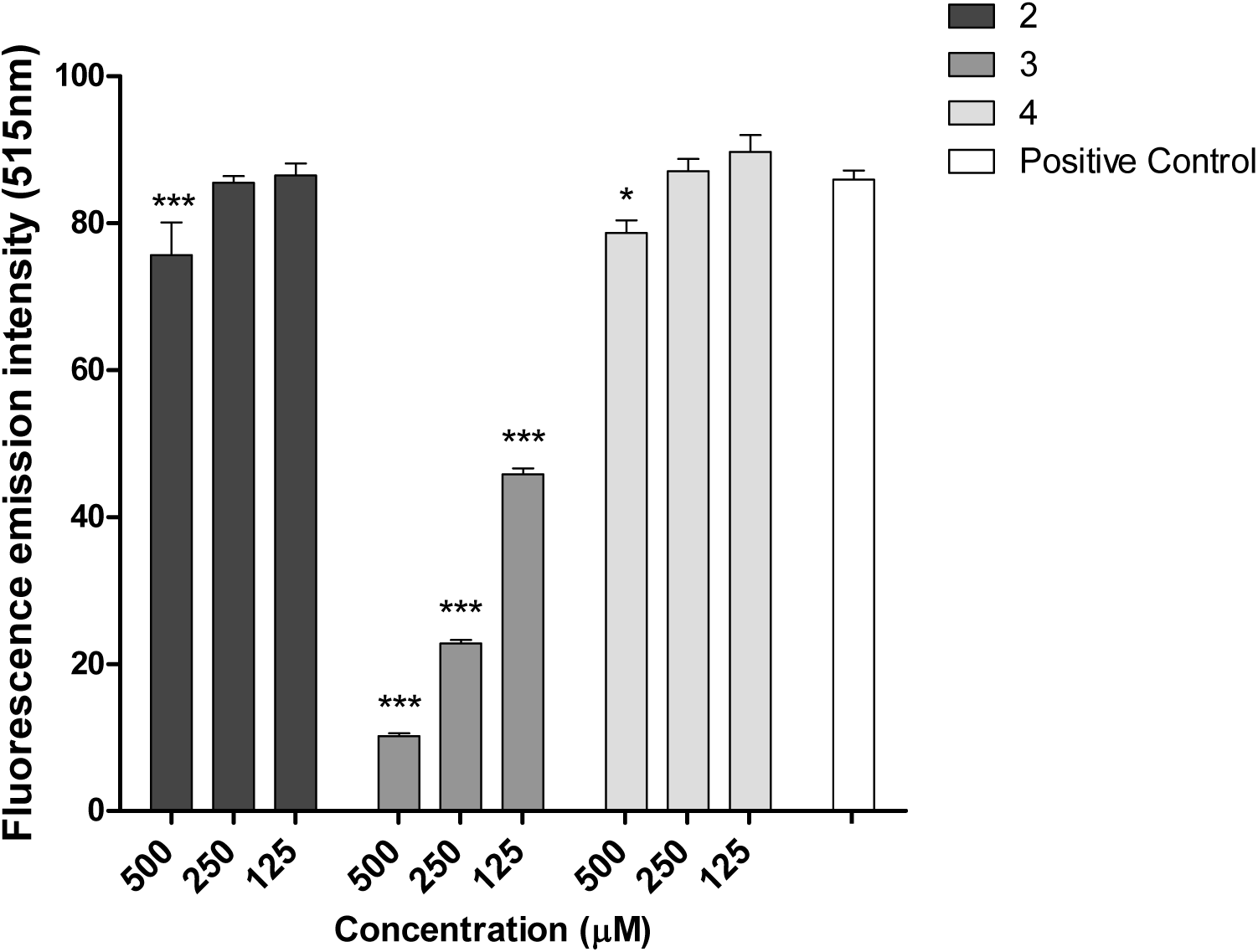
Inhibition of *B. asper* venom PLA₂ activity by ergosterol and ergostane-derived compounds using a liposome mix fluorescent substrate. Data are presented as RFU ± SEM (n = 3). *** indicates statistically significant differences compared with the positive control at *p* < 0.001, ** at *p* < 0.01.

**Figure 6.**
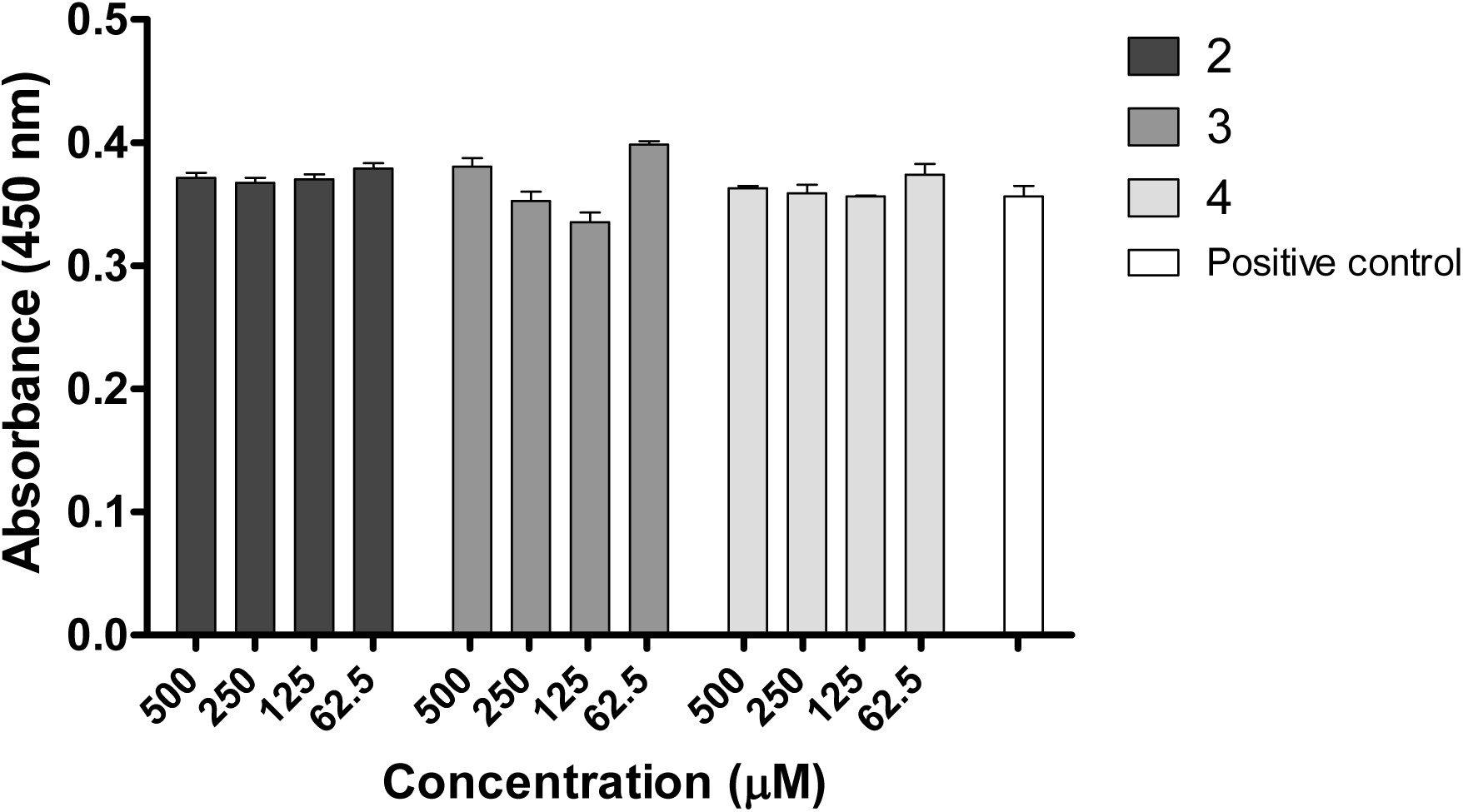
Inhibition of the proteolytic activity of *B. asper* venom by ergostane-derived compounds at λ = 450 nm. Data are presented as mean ± SEM (n = 3). *** indicates statistically significant differences compared with the positive control at p < 0.001, ** at p < 0.01, and * at p < 0.05.

SVMPs are zinc-dependent enzymes that play a central role in the pathophysiology of envenomation, particularly in the induction of local and systemic hemorrhage by degrading basement membrane components, including type IV collagen, laminin, and fibronectin (FN). This disruption compromises capillary integrity, leading to extravasation(43,44). As the effects of SVMPs are predominantly zinc-dependent, inhibitors that interact with Zn^2+^ to abrogate their catalytic activity have been investigated. Currently, two classes of SVMP inhibitors have been studied, both with a distinct mode of action: peptidomimetic inhibitors and metal chelators(43).

The evaluated compounds exhibited inhibitory activity against phospholipase A₂ (PLA₂) and serine proteinases, suggesting their selectivity for these toxin families. This specificity may result from differences in the enzyme structure and catalytic mechanisms. Ergosterol-derived compounds may preferentially interact with hydrophobic sites in PLA₂s and serine proteinases but lack the functional groups required for effective Zn^2+^ chelation or binding to the metalloproteinase active site. From a therapeutic perspective, such selectivity is advantageous because it allows targeting of specific pathological effects of envenomation, such as coagulopathy or myonecrosis, while minimizing off-target interactions. However, the inability to inhibit SVMPs highlights a limitation, given their major role in hemorrhagic damage, and suggests that combination strategies or structural modifications may be needed to achieve broader efficacy.

### 3.7. Molecular modeling inhibition of ergostane derivatives

To comprehensively understand the interactions between the studied compounds and their inhibitory properties, both target proteins were analyzed experimentally and computationally. Initially, molecular docking was performed to identify the most favorable binding orientations, which were subsequently used as starting structures for molecular dynamics (MD) simulations of the selected compounds. In this study, compound **2** was evaluated in complex with phospholipase A_2_ (PLA_2_), whereas compounds **3** and **4** were analyzed using serine venom snake protease (SVSP) as the receptor. We studied the interaction of a PLA_2_ dimer, assuming it would inhibit protein-membrane coupling (45).

For compound **2,** the optimal docking pose indicated that binding occurred within a hydrophobic pocket, where the side-chain scaffold of the compound was accommodated (Figure 7A). This pocket is formed by residues Leu3, Phe8, Pro18, Phe141, and Pro142, consistent with the binding sites reported for known PLA_2_ inhibitors co-crystallized with homologous enzymes from other species(46). However, the interaction of compounds **3** and **4** possesses a strong interaction with Lys60 through hydrogen bonds and the stability of the cationic-parallel stacking with the triazoline-dienone fragment, maintaining the hydrophobic interactions.

**Figure 7.**
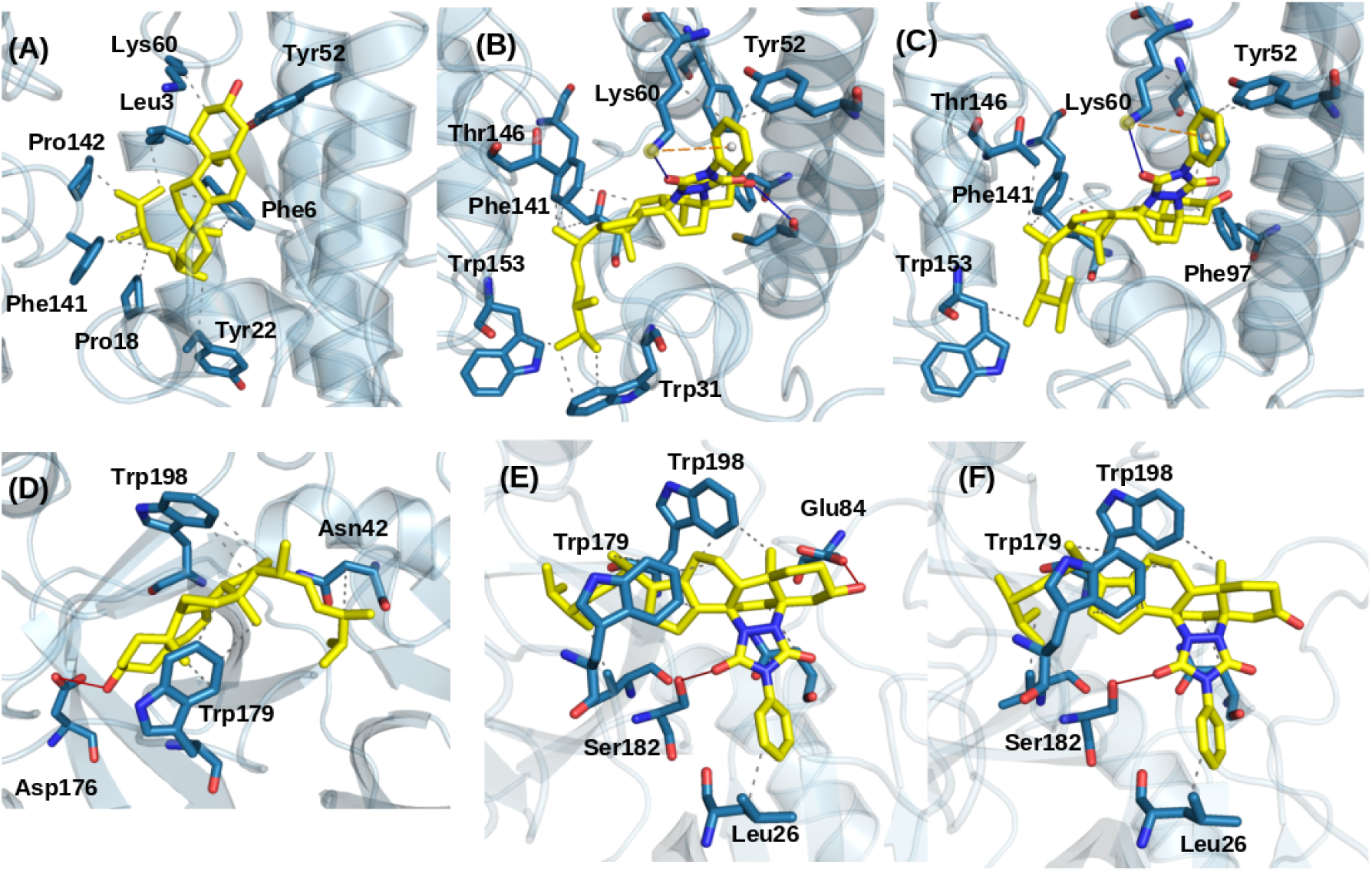
Molecular docking pose: (A-C) compound **2,3 and 4** in complex with PLA_2,_ (D-F) compound **2,3 and 4** in complex with SVSP.

The binding modes of compounds **3** and **4** in the SVSP complex were also examined in detail. Structural differences between the ergostane scaffold and the triazoline-dienone moiety were considered, as these features may influence the ligand behavior within the binding site. The docking results revealed that hydrophobic residues, particularly Trp179 and Trp198, play key roles in stabilizing the ergostane scaffold. Additionally, one of the carbonyl groups in the triazoline-dienone moiety adopts a preferred orientation that contributes to stabilizing the ligand–protein complex. In both cases, molecules **3-4** form hydrogen bonds with Ser182 (Figures 7E and 7F).

The stability and equilibration of the MD simulations were assessed by calculating the root-mean-square deviation (RMSD) of the backbone atoms relative to their initial structures. The RMSD profiles for all ligand–receptor complexes derived from the MD trajectories are presented in Figure S6 (Supplementary Data).

To evaluate the structural flexibility of ligand-receptor interactions, the Root-Mean-Square Fluctuations (RMSF), which measure the change in the compactness of the protein during the simulation, are typically utilized for equilibrated MD trajectories. In the RMSF analysis of residues (Figure 8), none of the complexes exhibited RMSF values exceeding 3 Å, except for residues located in the linker regions. Importantly, these fluctuations did not compromise ligand stability at the binding sites.

**Figure 8.**
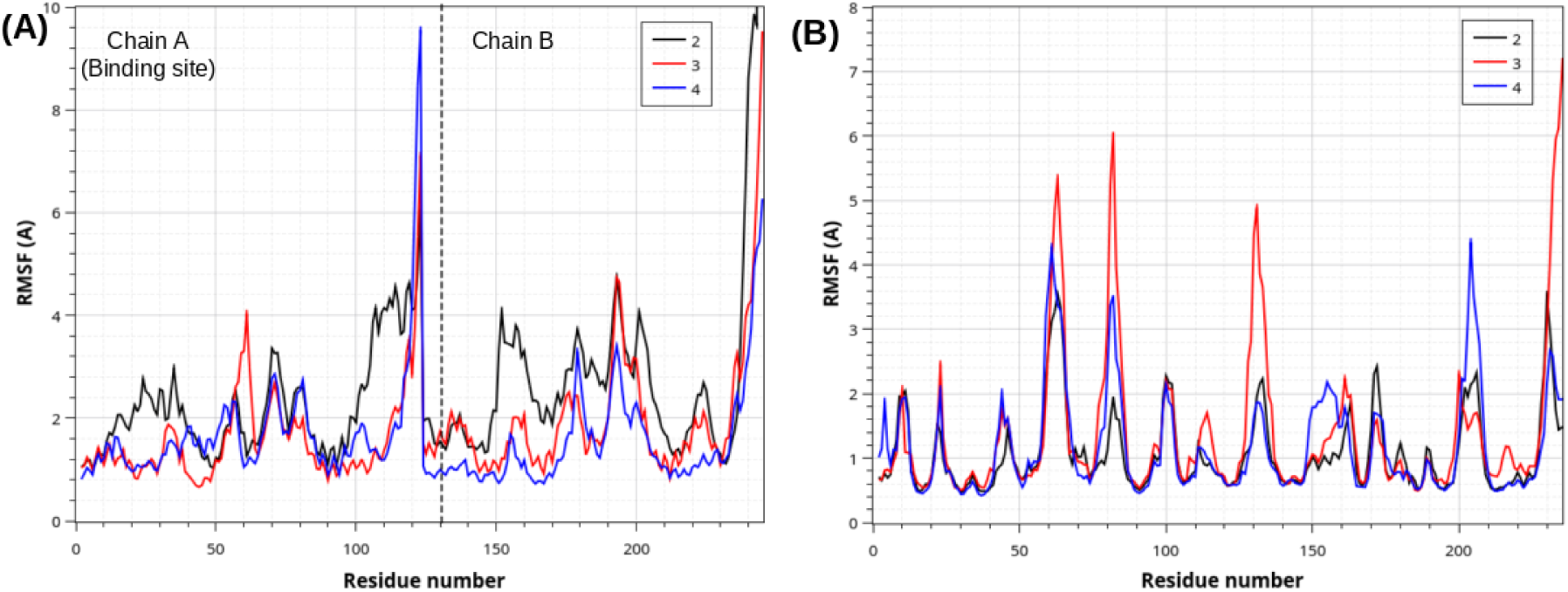
RMSF curve of the simulated system during MD simulation. (a) Compounds **2-4** in complex with PLA_2._ (b) Compounds **2-4** in complex with SVSP.

**Figure 8.**
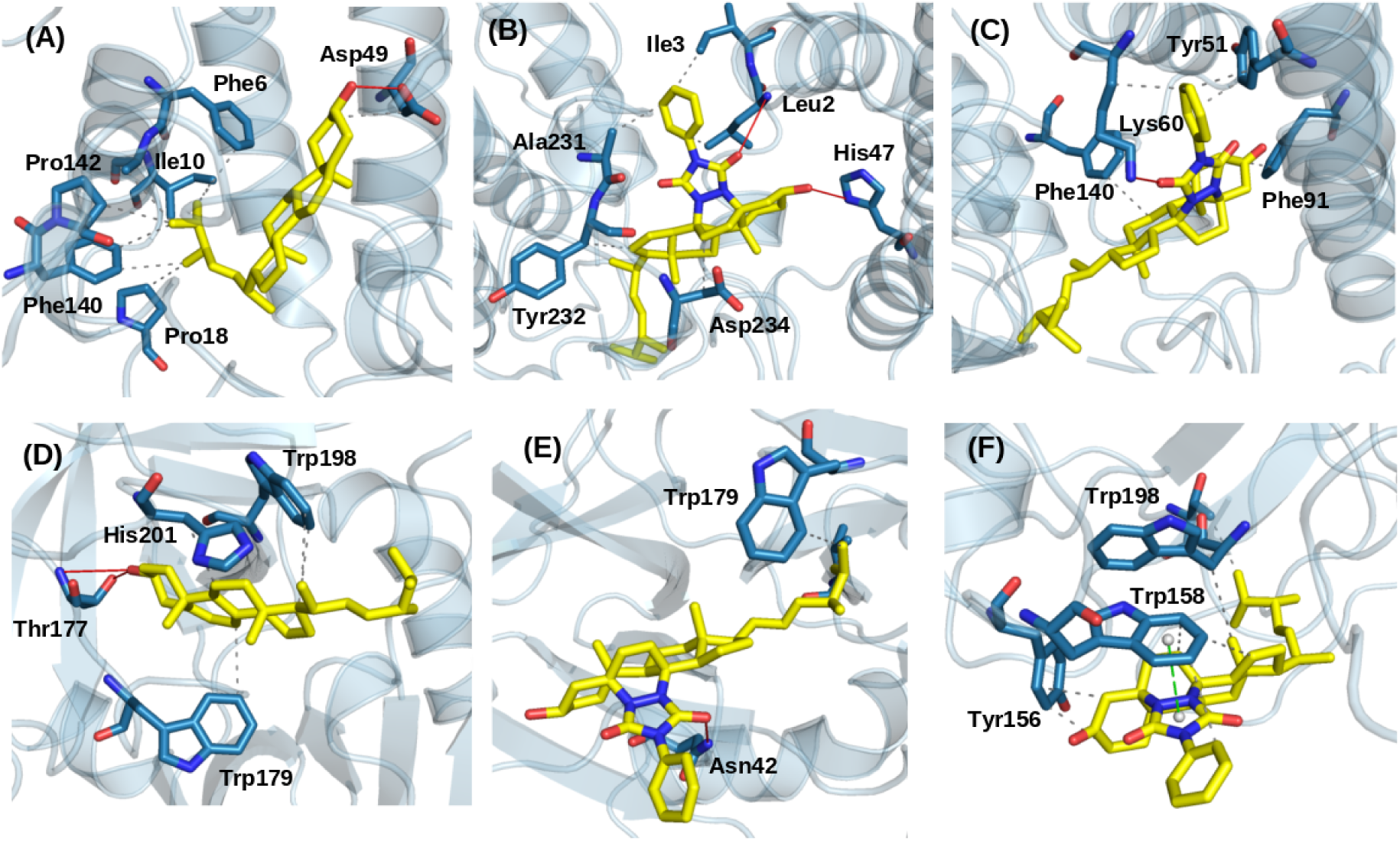
Representative cluster results of MD: (A-C) Compounds **2,3, and 4** in complex with PLA_2_ (D-F) compounds **2, 3, and 4** in complex with SVSP.

For the interaction between compound **2** and PLA_2_, the fluctuation between residues 20-40 is associated with a loop involving the ligand-receptor interaction region, although these variations remain below 3 Å. The highest fluctuations occurred between residues 100 and 120, which form solvent-exposed terminal loops. Furthermore, fluctuations corresponding to protein-protein interactions were observed in the region between residues 140 and 160, indicating that this interaction with the ligand was favored. Note that the fluctuation between residues 190 and 220 corresponds to a loop that is structurally analogous to the region spanning residues 100–120. Comparative RMSF analyses of compounds **3** and **4** bounds to SVSP revealed distinct fluctuation profiles, reflecting differences in their binding modes. Residues 60–70 correspond to a solvent-exposed loop, whereas the region around residue 140, which is also solvent-accessible, is located near the ligand-binding site. Notably, increased fluctuations were observed around residue 80 in the compound **3** complex, suggesting reduced stabilization relative to that of compound **4**. This observation indicates that compound **2** and **4** may confer greater stability to the ligand–receptor complex.

The simulation trajectory was clustered to extract representative conformations using the TTClust software[36]. Figures S7-S11 (supplementary data) show the first and last cluster representatives of the ligand-receptor complexes, along with the mode of interaction in an enlarged portion of the image. The figure shows the most representative clusters and their characteristic interactions. For compound **2** in complex with PLA_2_, hydrophobic interactions were the predominant interactions, as demonstrated by the overlay of the cluster analysis and the most populated cluster. In addition, a hydrogen bond is formed with residue Asp49. For compounds **3** and **4** complexed to PLA_2_, the triazoline-dienone moiety engages in hydrophobic interactions with residues Trp156, Trp179, and Trp198. Notably, the interaction between compound **4** and Trp156 involves π–π stacking, which likely contributes to the increased stability of this complex.

In the SVSP complexes, the binding mode of compound **2** shows a hydrogen bond with Thr177, along with consistent hydrophobic contacts with Trp179 and Trp198 throughout the molecular dynamic simulation. On the other hand, compounds **3** and **4** exhibited hydrophobic interactions between the ergostane scaffold and Trp residues, particularly Trp158, Trp179, and Trp198. Additionally, compound **4** displayed π–π stacking interactions between the triazoline ring and Trp158. These interactions, together with consistent hydrophobic contacts involving Trp179 and Trp198, suggest a preferred binding mode for ergostane derivatives. Notably, the compound **4**–SVSP complex showed significant fluctuations proximal to residues 195-210.

The binding free energy for the inhibitors in each system was calculated using the MM-GBSA approach. The binding energy of the compound **2**–PLA_2_ complex was −51.75 kcal/mol, while that of the compound 2–SVSP complex was −32.24 kcal/mol. The binding energies of compounds **3** and **4** in complex with PLA_2_ were determined to be −54.55 and −52.07 kcal/mol, respectively. For the SVSP complexes, the binding energies of compounds **3** and 4 were −31.86 and −34.0 kcal/mol, respectively (Table 1).

**Table 1.** Binding energy values and individual component energies were calculated using the MM-GBSA method for the PLA_2_ and SVSP complexes (values in kcal/mol).

| Enzym<br>e | Compound<br>d | $\Delta E_{\text{vdW}}$ | $\Delta E_{\text{ele}}$ | $\Delta E_{\text{GB}}$ | $\Delta E_{\text{surf}}$ | $\Delta G_{\text{bind}}$ |
| --- | --- | --- | --- | --- | --- | --- |
| PLA<sub>2</sub> | **2** | -54.95 ± 3.70 | -11.18 ± 6.47 | 20.96 ± 6.39 | -6.58 ± 0.55 | -51.75 ± 4.11 |
|  | **3** | -66.69 ± 4.24 | -15.67 ± 8.29 | 34.32 ± 7.31 | -6.50 ± 0.52 | -54.55 ± 5.40 |
|  | **4** | -63.66 ± 3.99 | -9.49 ± 6.11 | 29.36 ± 5.81 | -8.27 ± 0.34 | -52.07 ± 4.72 |
| SVSP | **2** | -36.77 ± 5.99 | -7.38 ± 3.60 | 15.82 ± 2.61 | -3.90 ± 0.57 | -32.24 ± 6.68 |
|  | **3** | -41.17 ± 5.24 | -4.99 ± 5.03 | 19.08 ± 5.12 | -4.77 ± 0.58 | -31.86 ± 5.29 |
|  | **4** | -43.96 ± 5.53 | -3.38 ± 6.36 | 17.99 ± 4.99 | -4.66 ± 0.56 | -34.02 ± 5.00 |

Energies: $\Delta E_{\text{vdW}}$ : van der Waals energy; $\Delta E_{\text{ele}}$ : Electrostatic energy; $\Delta E_{\text{GB}}$ : electrostatic contribution to the solvation free energy; $\Delta E_{\text{polar}}$ : nonpolar solvation energy; $\Delta G_{\text{bind}}$ : Estimated binding free energy.

These interaction energies are consistent with the experimental results, showing a strong correlation with the observed biological activities. For PLA_2_, the results reflect the activity trend of compound **3**, as well as the relationship between compounds **2** and **4**, in agreement with the inhibition data obtained from the fluorescence-based assay. At the same time, the relatively close values suggest that all the studied compounds have good affinity, supporting their role in inhibiting esterase activity associated with PLA_2_. In contrast, the amidolytic activity, as reflected by the inhibition of the SVSP enzyme, was consistent with the experimental observations.

In addition, van der Waals (EvdW) contributions were the largest contributors to the interaction energy across all compounds because of the hydrophobic nature of the ergostane scaffold, which appears to be the most significant factor governing ligand–receptor interactions.

In contrast, the electrostatic energy (E_ele_) also contributed significantly to the binding affinity, underscoring its importance in complex formation. In contrast, polar solvation energies (E_GB_) were unfavorable, likely due to the size of the binding pocket and the partial solvent exposure of the hydrophobic ligands. However, for the molecules, the results suggest that complex formation is favored by the intermolecular electrostatic and van der Waals interactions, as well as the nonpolar component of the solvation free energy repulsive term (hydrophobic/cavity) energies. In contrast, nonpolar solvation free energies were favorable for ligand binding.

Figure 9 illustrates the per-residue binding energy decomposition obtained from MM/GBSA calculations for representative compound–protein systems. For the compound **2**–PLA2 complex, the residues contributing most significantly to the binding energy are located within the ligand interaction region. These contributions are consistent with the compound’s high affinity for chain A, as also reflected in the RMSF analysis. In this context, Leu3, Phe6, Ile445, and Phe140 are the main contributors to ligand–receptor interactions. For compound **3**, the most stabilizing interactions involve residues Leu3, Tyr145, and Ala232. In the case of compound **4**, greater stabilization is observed through interactions with residues Lys60, Ala91, Phe96, Phe140, and Trp152. This suggests a more stable complex, in agreement with the calculated interaction energy values.

**Figure 9.**
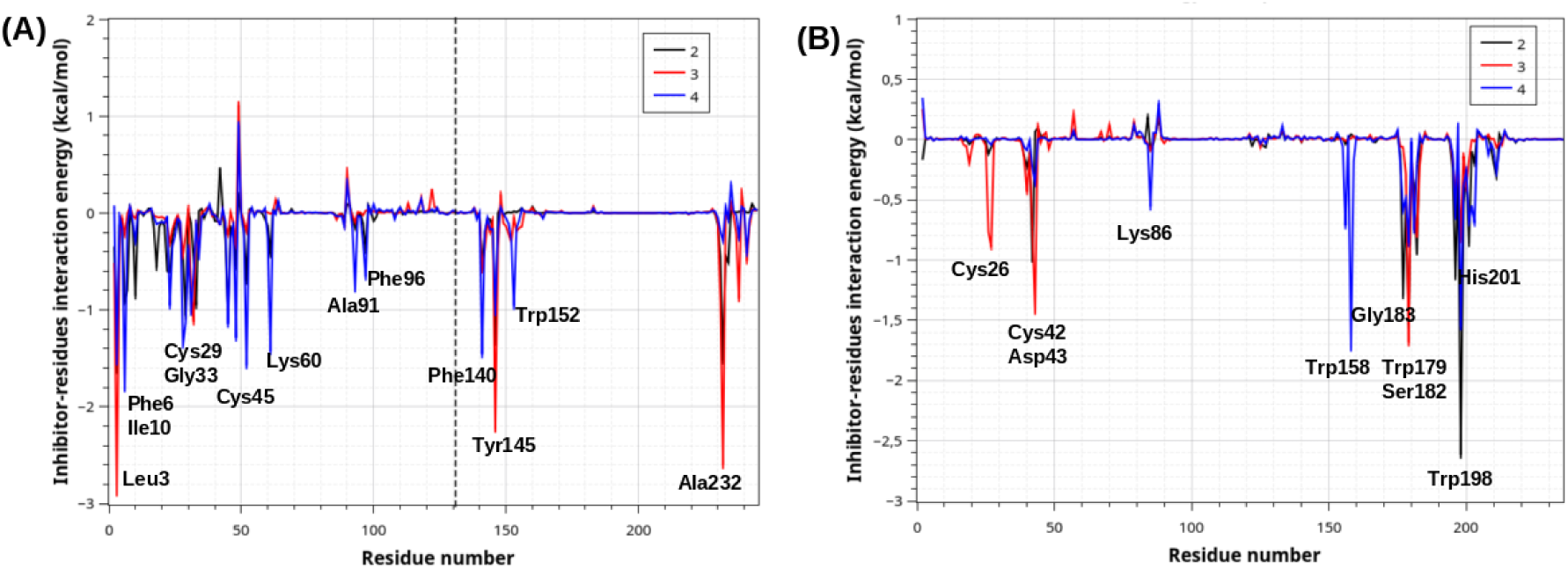
Binding energy decomposition plots for the systems: (A) compound **2-4** with PLA_2_ and (B) compounds **2-4** with SVSP.

For the SVSP complexes, the interaction patterns differed owing to the distinct ligand orientations during MD simulations. For compound **2** in complex with SVSP, the greatest contribution to the interaction energy would be generated by the interaction with Trp198, due to the hydrophobic interaction. In the **3**–SVSP complex, hydrophobic interactions with Trp178 and hydrogen bonding with Ser128 were the primary contributors to the activation energy. Conversely, in the 4–SVSP complex, π–π stacking with Trp158 and hydrophobic interactions with Trp198 dominated the binding energetics, which is in agreement with the clustering analysis. Overall, these computational findings provide a mechanistic explanation for the experimentally observed activities and support the potential of ergostane-type compounds as promising scaffolds for the development of inhibitors targeting PLA_2_ and SVSP enzymes.

## 4. Conclusion

The present study provides an integrated experimental and computational evaluation of ergosterol-derived compounds as potential inhibitors of key enzymatic components of *Bothrops asper* venom. Together, the *in vitro* assays and molecular modeling analyses demonstrated that these compounds inhibited phospholipases A₂ (PLA₂) and serine venom snake proteases (SVSP), two enzymes that play central roles in the local and systemic effects of envenomation. Importantly, the absence of inhibitory activity against metalloproteinases suggests a degree of selectivity in the mechanism of action. Docking and molecular dynamics simulations provide mechanistic insights into the molecular basis of enzyme inhibition. For PLA₂, compound **2,3** and **4** preferentially bound to a conserved hydrophobic pocket involved in protein–membrane interactions, supporting the hypothesis that dimer targeting may interfere with enzymatic function. The stability of this interaction was reinforced by MD analyses, which revealed limited structural fluctuations and strong hydrophobic contributions to the binding free energy. For SVSP, compounds **2**, **3,** and **4** exhibited distinct binding modes, primarily driven by hydrophobic interactions with Trp-rich regions and, for compounds **3** and **4**, by additional π–π stacking interactions that enhanced complex stability. These findings are consistent with the MM-GBSA calculations, which highlighted van der Waals and electrostatic interactions as the main energetic contributors to ligand binding.

Notably, discrepancies between the predicted binding affinities and biological outcomes highlight the complexity of venom enzyme modulation. The observation that a compound with a comparatively lower theoretical affinity displayed a clear dose-dependent anticoagulant effect suggests that these ergosterol derivatives may act through alternative or allosteric mechanisms that are not fully captured by classical docking approaches. This emphasizes the need to combine computational predictions with experimental validation when characterizing these venom inhibitors. Despite the limitations related to compound solubility and partial inhibition, this study provides novel evidence supporting the antivenom potential of ergosterol-based scaffolds. Overall, the combined experimental and computational results indicate that these compounds represent promising lead structures for the development of complementary therapies to conventional antivenoms. Further studies, including detailed mechanistic investigations and *in vivo* evaluations, are essential to fully elucidate their mode of action and assess their therapeutic efficacy and safety in preclinical models.

## Acknowledged

MB thanks the Fondecyt Project N° 3220756. YAR-N. We thank the Fondecyt project N° 11241068 and UNAB for financial support (project Dl-05.23/REG). LMP acknowledges the support of the CODI project 2020-33639 (CIFAL-321) and CIFAL-377.

## Author contributions

MB, YAR-N, EP-C conceived and designed the chemical experiments. ATR, JMA and LPR perform venom inhibition experiments. CJG, JS-D analyzed the Docking and MD tests. MB, YAR-N, ATR, JMA, LPR, EP-C, CJG, JS-D wrote the article.

## References

1. World Health Organization. https://www.who.int/news-room/fact-sheets/detail/snakebite-envenoming. 2023. Snakebite envenoming.

2. Instituto Nacional de Salud - Dirección de Vigilancia y Análisis del Riesgo en Salud Pública. Protocolo de vigilancia en salud pública. https://www.ins.gov.co/buscador-eventos/Lineamientos/Pro_AO_Venenosos_2024.pdf. 2024. Accidente Ofídico - Accidentes por otros animales venenosos.

3. Instituto Nacional de Salud. https://app.powerbi.com/view?r=eyJrIjoiMjI2MjAzYjYtMzE5YS00MmM1LTk1ZGEtOTUxYjFiNjlhZjNmIiwidCI6ImE2MmQ2YzdiLTlmNTktNDQ2OS05MzU5LTM1MzcxNDc1OTRiYiIsImMiOjR9. 2025. Accidente Ofídico.

4. Otero-Patiño R. Snake Bites in Colombia. In. 2018. p. 3–50. doi:10.1007/978-94-017-7438-3_41

5. Pereañez JA, Preciado LM, Rey-Suárez P. Knowledge about Snake Venoms and Toxins from Colombia: A Systematic Review. Toxins (Basel). 2023 Nov 15;15(11):658. doi:10.3390/toxins15110658

6. World Health Organization. https://www.who.int/teams/control-of-neglected-tropical-diseases/snakebite-envenoming/antivenoms. 2024. Antivenoms.

7. Maria Gutierrez J, Leon G, Lomonte B, Angulo Y. Antivenoms for Snakebite Envenomings. Inflamm Allergy Drug Targets. 2011 Oct 1;10(5):369–80. doi:10.2174/187152811797200669

8. Calvete JJ, Juárez P, Sanz L. Snake venomics. Strategy and applications. Journal of Mass Spectrometry. 2007 Nov 10;42(11):1405–14. doi:10.1002/jms.1242

9. Lomonte B, León G, Angulo Y, Rucavado A, Núñez V. Neutralization of Bothrops asper venom by antibodies, natural products and synthetic drugs: Contributions to understanding snakebite envenomings and their treatment. Toxicon. 2009 Dec;54(7):1012–28. doi:10.1016/j.toxicon.2009.03.015

10. Gao JM, Wang M, Liu LP, Wei GH, Zhang AL, Draghici C, et al. Ergosterol peroxides as phospholipase A2 inhibitors from the fungus Lactarius hatsudake. Phytomedicine. 2007 Dec;14(12):821–4. doi:10.1016/j.phymed.2006.12.006

11. Oliveira AL, Viegas MF, da Silva SL, Soares AM, Ramos MJ, Fernandes PA. The chemistry of snake venom and its medicinal potential. Nat Rev Chem. 2022 Jun 10;6(7):451–69. doi:10.1038/s41570-022-00393-7

12. Ghorbani-Choghamarani A, Chenani Z, Mallakpour S. Supported nitric acid on silica gel and polyvinyl pyrrolidone (PVP) as an efficient oxidizing agent for the oxidation of urazoles and bis-urazoles. Synth Commun. 2009 Jan;39(23):4264–70. doi:10.1080/00397910902898619

13. Theakston RD, Reid HA. Development of simple standard assay procedures for the characterization of snake venom. Bull World Health Organ. 1983;61(6):949–56. PubMed PMID: 6609011.

14. Theakston RD, Reid HA. Development of simple standard assay procedures for the characterization of snake venom. Bull World Health Organ. 1983;61(6):949–56. PubMed PMID: 6609011.

15. Erlanger BF, Kokowsky N, Cohen W. The preparation and properties of two new chromogenic substrates of trypsin. Arch Biochem Biophys. 1961 Nov;95(2):271–8. doi:10.1016/0003-9861(61)90145-X

16. Wang WJ, Shih CH, Huang TF. A novel P-I class metalloproteinase with broad substrate-cleaving activity, agkislysin, from Agkistrodon acutus venom. Biochem Biophys Res Commun. 2004 Nov;324(1):224–30. doi:10.1016/j.bbrc.2004.09.031

17. Pereañez J. JA, JSL, QJC, NV, FM, & RY. Inhibición de las actividades proteolítica, coagulante y hemolítica indirecta inducidas por el veneno de Bothrops asper por extractos etanólicos de tres especies de heliconias. Vitae. 2009;15(1).

18. Holzer M, Mackessy SP. An aqueous endpoint assay of snake venom phospholipase A2. Toxicon. 1996 Oct;34(10):1149–55. doi:10.1016/0041-0101(96)00057-8

19. Bateman A, Martin MJ, Orchard S, Magrane M, Adesina A, Ahmad S, et al. UniProt: the Universal Protein Knowledgebase in 2025. Nucleic Acids Res. 2025 Jan 6;53(D1):D609–17. doi:10.1093/nar/gkae1010

20. Laskowski RA, MacArthur MW, Moss DS, Thornton JM. PROCHECK: a program to check the stereochemical quality of protein structures. J Appl Crystallogr. 1993 Apr 1;26(2):283–91. doi:10.1107/S0021889892009944

21. Morris GM, Huey R, Lindstrom W, Sanner MF, Belew RK, Goodsell DS, et al. AutoDock4 and AutoDockTools4: Automated docking with selective receptor flexibility. J Comput Chem. 2009 Dec;30(16):2785–91. doi:10.1002/jcc.21256

22. Schrödinger LLC. The PyMOL Molecular Graphics System, Version 2.5. 2024 Nov.

23. Trott O, Olson AJ. AutoDock Vina: Improving the speed and accuracy of docking with a new scoring function, efficient optimization, and multithreading. J Comput Chem. 2009;NA-NA. doi:10.1002/jcc.21334

24. Schake P, Bolz SN, Linnemann K, Schroeder M. PLIP 2025: introducing protein–protein interactions to the protein–ligand interaction profiler. Nucleic Acids Res. 2025 Jul 7;53(W1):W463–5. doi:10.1093/nar/gkaf361

25. Case DA, Aktulga HM, Belfon K, Ben-Shalom IY, Berryman JT,, Brozell SR,. Amber 2023. University of California, San Francisco; 2023.

26. Kagami L, Wilter A, Diaz A, Vranken W. The ACPYPE web server for small-molecule MD topology generation. Bioinformatics. 2023 Jun 1;39(6). doi:10.1093/bioinformatics/btad350

27. Sousa da Silva AW, Vranken WF. ACPYPE - AnteChamber PYthon Parser interfacE. BMC Res Notes. 2012 Dec 23;5(1):367. doi:10.1186/1756-0500-5-367

28. Tubiana T, Carvaillo JC, Boulard Y, Bressanelli S. TTClust: A Versatile Molecular Simulation Trajectory Clustering Program with Graphical Summaries. J Chem Inf Model. 2018 Nov 26;58(11):2178–82. doi:10.1021/acs.jcim.8b00512

29. Miller BR, McGee TD, Swails JM, Homeyer N, Gohlke H, Roitberg AE. MMPBSA.py : An Efficient Program for End-State Free Energy Calculations. J Chem Theory Comput. 2012 Sep 11;8(9):3314–21. doi:10.1021/ct300418h

30. Humphrey W, Dalke A, Schulten K. VMD: Visual molecular dynamics. J Mol Graph. 1996 Feb;14(1):33–8. doi:10.1016/0263-7855(96)00018-5

31. Ghorbani-Choghamarani A, Zolfigol MA, Hajjami M, Mallakpour S. Metal-free catalytic oxidation of urazoles under mild and heterogeneous conditions via combination of ammonium nitrate and catalytic amounts of silica sulfuric acid. Journal of the Iranian Chemical Society. 2010 Dec;7(4):834–9. doi:10.1007/BF03246076

32. Cookson RC, Gilani SSH, Stevens IDR. Diels–Alder reactions of 4-phenyl-1,2,4-triazoline-3,5-dione. J Chem Soc C. 1967;0(0):1905–9. doi:10.1039/J39670001905

33. Zolfigol MA, Zebarjadian MH, Chehardoli G, Mallakpour SE, Shamsipur M. An efficient method for the oxidation of urazoles with [NO+·crown·H(NO3)2−]. Tetrahedron. 2001 Feb;57(8):1627–9. doi:10.1016/S0040-4020(00)01150-9

34. Bourgeois F, Höller U, Netscher T. Synthesis of trifold-labeled versatile reagent [3,5- ^13^ C _2_, 4-^15^N]4-phenyl-1,2,4-triazoline-3,5-dione. J Labelled Comp Radiopharm. 2023 Dec 20;66(14):461–6. doi:10.1002/jlcr.4067

35. Mora-Obando D, Salazar-Valenzuela D, Pla D, Lomonte B, Guerrero-Vargas JA, Ayerbe S, et al. Venom variation in Bothrops asper lineages from North-Western South America. J Proteomics. 2020 Oct;229:103945. doi:10.1016/j.jprot.2020.103945

36. Mora-Obando D, Lomonte B, Pla D, Guerrero-Vargas JA, Ayerbe-González S, Gutiérrez JM, et al. Half a century of research on Bothrops asper venom variation: biological and biomedical implications. Toxicon. 2023 Jan;221:106983. doi:10.1016/j.toxicon.2022.106983

37. Serrano SMT. The long road of research on snake venom serine proteinases. Toxicon. 2013 Feb;62:19–26. doi:10.1016/j.toxicon.2012.09.003

38. Kini RM. Anticoagulant proteins from snake venoms: structure, function and mechanism. Biochemical Journal. 2006 Aug 1;397(3):377–87. doi:10.1042/BJ20060302

39. Tonello F, Rigoni M. Cellular Mechanisms of Action of Snake Phospholipase A2 Toxins. In: Snake Venoms. Dordrecht: Springer Netherlands; 2017. p. 49–65. doi:10.1007/978-94-007-6410-1_26

40. Gao JM, Wang M, Liu LP, Wei GH, Zhang AL, Draghici C, et al. Ergosterol peroxides as phospholipase A2 inhibitors from the fungus *Lactarius hatsudake*. Phytomedicine. 2007 Dec;14(12):821–4. doi:10.1016/j.phymed.2006.12.006

41. Jørgensen K, Davidsen J, Mouritsen OG. Biophysical mechanisms of phospholipase A2 activation and their use in liposome-based drug delivery. FEBS Lett. 2002 Oct 30;531(1):23–7. doi:10.1016/S0014-5793(02)03408-7

42. Burke JE, Dennis EA. Phospholipase A2 structure/function, mechanism, and signaling. J Lipid Res. 2009 Apr;50:S237–42. doi:10.1194/jlr.R800033-JLR200

43. Fox JW, Serrano SMT. Structural considerations of the snake venom metalloproteinases, key members of the M12 reprolysin family of metalloproteinases. Toxicon. 2005 Jun;45(8):969–85. doi:10.1016/j.toxicon.2005.02.012

44. Gutiérrez JM, Rucavado A, Escalante T, Díaz C. Hemorrhage induced by snake venom metalloproteinases: biochemical and biophysical mechanisms involved in microvessel damage. Toxicon. 2005 Jun;45(8):997–1011. doi:10.1016/j.toxicon.2005.02.029

45. Salvador GHM, dos Santos JI, Lomonte B, Fontes MRM. Crystal structure of a phospholipase A2 from Bothrops asper venom: Insights into a new putative “myotoxic cluster.” Biochimie. 2017 Feb;133:95–102. doi:10.1016/j.biochi.2016.12.015

46. Crous C, Petzer A, Petzer JP. Interactions of small molecule inhibitors with secreted phospholipase A _2_ : A review of the structural data. Chem Biol Drug Des. 2024 Feb 29;103(2). doi:10.1111/cbdd.14466

